# Integrative systems biology framework discovers common gene regulatory signatures in multiple mechanistically distinct inflammatory skin diseases

**DOI:** 10.1101/2023.10.24.563852

**Authors:** Bharat Mishra, Mohammad Athar, M. Shahid Mukhtar

## Abstract

**Aim:** More than 20% of the population across the world is affected by non-communicable inflammatory skin diseases including psoriasis, atopic dermatitis, hidradenitis suppurativa, rosacea, etc. Many of these chronic diseases are painful and debilitating with limited effective therapeutic interventions. However, recent advances in psoriasis treatment have improved the effectiveness and provide better management of the disease. Similar effective treatments are also needed for other chronic skin disorders regulated molecular pathogenesis. This study aims to identify common regulatory pathways and master regulators that regulate molecular pathogenesis.

**Methods:** We designed an integrative systems biology framework to identify the significant regulators across several inflammatory skin diseases. We exploited the gene co-expression-, and protein-protein interaction-based networks to identify shared genes and pro components in different diseases with relevant functional implications. Further, we utilized network analytics to unravel the high-priority proteins, that play a crucial role in disease pathogenesis.

**Results:** With conventional transcriptome analysis we identified 55 shared genes, which are enriched in several immune-associated pathways in eight inflammatory skin diseases. Our correlation-based co-expression networks identified the core co-expression networks and the integrative multi-omics interactomes providing core interactions across four of the eight diseases. Next, by implementing network analytics, we unraveled 55 high-value proteins as significant regulators in molecular pathogenesis. We believe that these significant regulators should be explored with critical experimental approaches to identify the putative drug targets for more effective treatments. As an example, we identified IKZF1 as a shared significant master regulator in three inflammatory skin diseases, which can serve as a putative drug target with known disease-derived molecules for developing efficacious combinatorial treatments for hidradenitis suppurativa, atopic dermatitis, and rosacea.

**Conclusion:** The conventional transcriptome results identify huge complexity among inflammatory skin diseases, which demands a systems biology perspective for comprehensive understanding. The proposed framework is very modular, which can indicate a significant path of molecular mechanism-based drug development from complex transcriptomics data and other multi-omics data.

## INTRODUCTION

Nearly 25% of the world’s population is affected by non-communicable chronic inflammatory diseases^1^. Some of the inflammatory skin diseases include acne, atopic dermatitis (AD), actinic keratoses (AK), psoriasis (PS), hidradenitis suppurativa (HS), and three types of rosacea (RS) share specific immune regulator activities with psoriasis^1–3^. Most of these diseases manifest autoimmune signatures on a temporal basis. Psoriasis is one of the best-described inflammatory skin diseases that affects approximately 1-3% of global adults^4^. Psoriasis patients manifest comorbidities such as diabetes, metabolic disorders, and severe cardiovascular disease^5^. Previous reports suggested that keratinocytes are major drivers of psoriasis in the skin epidermis, which was later expanded to the involvement of immune cells, specifically T helper 17 cells (Th17) are the key drivers of disease^6,7^. Eventually, targeting Th17 cell signaling and several interleukin cytokines including IL-23, IL-17A, and IL-17F have improved long-term remissions by 85-100% in psoriasis patients^5,7^. Similarly, AD is an immune-mediated disease that is prevalent in approximately 25% of children and 10% of adults^8^. Likewise, HS is associated with inflammatory bowel disease and has a distinct anatomical etiology including the occurrence of inflammatory nodules, abscesses, and pus-draining sinus tracts at the places where skin rubs against each other like the underarm and groin^6^. The treatment regimens for AD and HS have very little effectiveness and cannot serve the majority of moderate to severe disease patients. Therefore, understanding the shared and unique genetic signatures, activated pathways, and regulators through innovative systems biology and integrative multi-omics can be to our advantage.

The omnipresence of bulk transcriptomics dataset introduces an extremely rewarding line of investigation to systems biologists for integration of transcriptomics to other omics layers including biological networks for a comprehensive and extrapolated exploration of some limited-studied diseases including immune-mediated diseases^9^. Network sciences have been applied to diverse biological domains to identify the associated biomolecules in disease, cancer, infections, and other stress conditions. In simple terms, biological networks are graphical representations containing nodes (genes/proteins) and edges (links) for a specific condition^10,11^. These networks tend to change their interacting partners in different scenarios including normal/healthy and disease response^12^. Network-based systems biology analytics has proven a great tool to unravel the structural properties of biological networks and highlight the significant contributors to disease pathogenesis and organismal development across different systems^9,13–18^. In this study, we designed an integrative systems biology framework to identify the significant regulators across several inflammatory skin diseases. We exploited the gene co-expression network (GCN)- and protein-protein interaction (PPI)-based networks to identify shared genes and protein components in eight skin diseases with relevant functional implications. Further, we identified 55 high-priority proteins (HPPs) with increased network indices, which are also associated with immune-mediated pathways. Finally, we explored the candidate drug-gene interactions to highlight some therapeutic-relevant target selection for putative treatment strategies. In summary, our integrative framework unravels the data-based regulatory signatures and activated molecular pathways across inflammatory skin diseases.

## METHODS

### Data acquisition

Publicly available transcriptome datasets were extracted from NCBI GEO in eight different inflammatory skin diseases including hidradenitis suppurativa (HS, GSE72702)^19^, rosacea (RS, GSE65914)^20^, atopic dermatitis (AD, GSE121212)^21^, contact dermatitis (CD, GSE6281)^22^, actinic keratoses (AK, GSE90643)^23^, Irritant contact dermatitis (ICD, GSE18206)^24^, Acne (GSE53795)^25^, and psoriasis (PS, GSE121212)^21^.

### Transcriptomics analysis

The differentially expressed genes (DEGs) analysis was performed on all eight disease datasets with the same threshold parameters ^26^. DESeq2 was used for RNA-Seq experiments and edgeR was used for microarray experiments.

### Co-expression network constriction

We performed the gene co-expression network (GCN) construction and analysis for all eight disease datasets through weighted gene co-expression network analysis WGCNA^27^. We achieved eight disease-specific networks ranging from 320 nodes with 264 edges in Acne to 5,465 nodes with 132,849 edges in PS. This analysis demonstrated the challenges in the handling of multivariate transcriptomes. Networks were visualized in Cytoscape^28^.

### Protein-protein interaction network construction and analysis

To establish the disease-specific protein-protein interaction network we fused the results from co-expression networks, DEGs, and the largest publicly available human protein-protein interaction network from the STRING database^29^. Finally, we had a collection of eight disease-specific protein-protein interaction networks ranging from 90 nodes with 90 edges in Acne to 3,622 nodes with 33,173 edges in PS. Next, we performed network centrality analyses on all four filtered networks to identify the most important players in each disease^9^. Specifically, we analyzed the networks by betweenness centrality (bottlenecks), degree (connections), eigenvalue (combined effect of degree and betweenness), information centrality (ability to pass stimuli information), and inner core proteins identification by weighted *k*-sell decomposition method^13^. Networks were visualized in Cytoscape^28^.

### Network exposibility analysis

To check the extent of coverage in a network and inspired by the COVID-19 exposibility scenario, we calculated the network exposibility by dividing the total number of nodes by the total connected components in the network. Based on the network exposibility threshold ≥ 10 to determine the reliability of eight PPI networks, we filtered four disease-specific interaction networks including HS, PS, AD, and RS for further analysis.

### High-priority protein identification

To identify the High-priority proteins (HPPs) or significant proteins for each PPI network we calculated the top 5% of centrality value nodes with the degree, betweenness, eigenvector, information centrality, and the inner layer proteins from the weighted *k*-shell decomposition method. Next, if these significant proteins are also significantly activated or inhibited regulators characterized by pathway analysis in each of the eight inflammatory skin diseases, then these proteins are termed HPPs or significant regulators. Additionally, we identified the most important regulators shared among four diseases and mapped them against their regulated pathways.

### Drug-gene interaction network

We extracted the HPPs interacting drugs from the publicly available databases DGIdb^30^. Only the interactions with listed publication information were included in this analysis. These drug-gene (HPPs) interactions can provide significant information about the potential therapeutic options for a disease or class of diseases. The drug-gene interaction network was visualized in Cytoscape^28^.

### Gene Ontology and Pathway Analysis

The canonical pathway and regulator analysis was performed by IPA with default parameters. The gene ontology and pathway analysis were performed through metascape^31^ with standard parameters.

### Significance analysis

The DEG analysis was performed with (log2*FC* ≥|1|; *FDR*<0.05) parameters. The canonical pathway analysis was performed with a significance test BH *P*-value < 0.05. The gene ontology analysis was performed with a significance test *P*-value < 0.05. The network power-law distribution fitness threshold is *r*^2^ ≥0.5. The network exposibility threshold was 10 for reliable networks. The correlation among values of different centralities is positive if *r*≥0.5. The significance of degree and betweenness among the inner and outer layers of four PPIs were tested by Mann-Whitney test for non-parametric distributed values. The significant regulator analysis through IPA was tested by BH *P*-value < 0.05.

## RESULTS

### A Network Science-based Framework to Prioritize Genes/Pathways from Transcriptome Datasets

To study the shared and unique genetic signatures, regulators, and prioritizing genes in different chronic inflammatory skin diseases, we implemented an integrative multi-omics framework utilizing the transcriptome datasets and extrapolating the network-based analysis pipeline. In this framework, 11 different steps are followed to identify the most appropriate drug-gene interactions (Figure 1). **1**. First, we extracted publicly available transcriptome datasets in eight different inflammatory skin diseases (namely: acne, atopic dermatitis (AD), actinic keratoses (AK), psoriasis (PS), hidradenitis suppurativa (HS), and three types of rosacea (RS)) as described in the methods. **2**. Subsequently, we performed the gene co-expression network construction and analysis for all eight disease datasets through WGCNA^27^. We achieved eight disease-specific networks ranging from 320 nodes with 264 edges in Acne to 5,465 nodes with 132,849 edges in PS. Which demonstrated the challenges in the handling of multivariate transcriptomes. **3**. The third step was to perform DEGs analysis on all eight disease datasets with the same threshold parameters^26^. We observed some similarities and differences in the expression profiles among eight diseases. **4**. To establish the disease-specific protein-protein interaction network, we next combined the results from co-expression networks, DEGs, and the largest publicly available human protein-protein interaction network from the STRING database^29^. Finally, we had a collection of eight disease-specific protein-protein interaction networks ranging from 90 nodes with 90 edges in Acne to 3,622 nodes with 33,173 edges in Psoriasis. **5**. Based on the network explosibility, a comparable number of nodes and interactions, as well as power-law distribution of eight networks, we selected four disease-specific interaction networks including atopic dermatitis (AD), psoriasis (PS), hidradenitis suppurativa (HS), and rosacea (RS) for further analysis. **6**. Afterwards, we performed network centrality analyses on all four filtered networks to identify the most important players in each disease^9^. Specifically, we analyzed the networks by betweenness centrality (bottlenecks), degree (hubs), eigenvalue (combined effect of degree and betweenness), information centrality (ability to pass stimuli information), and inner core proteins identification by the weighted *k*-sell decomposition method. **7**. With that, we obtained a collection of high-priority proteins (HPPs) in these four inflammatory skin diseases. When comparing the appearance of these HPPs in all four diseases, we identified shared and unique HPPs in each disease. **8**. Then, we mapped the HPPs in significantly activated/inhibited signaling pathways in these four skin diseases. Interestingly, we found some highly activated signaling pathways are shared among all four diseases. Additionally, we identified the most important regulators shared among four diseases and mapped them against their regulated pathways. **9**. Moving forward, we identified the HHPs interacting with drugs from the publicly available databases DGIdb^30^. These drug-gene (HPPs) interactions can provide significant information about the potential therapeutic options for a disease or class of diseases. **10**. From these data, we propose that these drugs can be repurposed for treating several inflammatory skin diseases either alone or in combination. **11**. All the predictive outcomes including the role of genes, pathways, HPPs, drug-HPP interactions, and other predictions are required to be validated experimentally for each disease condition. Taken together, we proposed a modular framework that can be applied to any polygenic chronic inflammatory disease to refine the regulator identification, pathway association, and drug-gene prediction for the alternative therapeutic targets in any immunogenetic-related disease.

**Figure 1.**
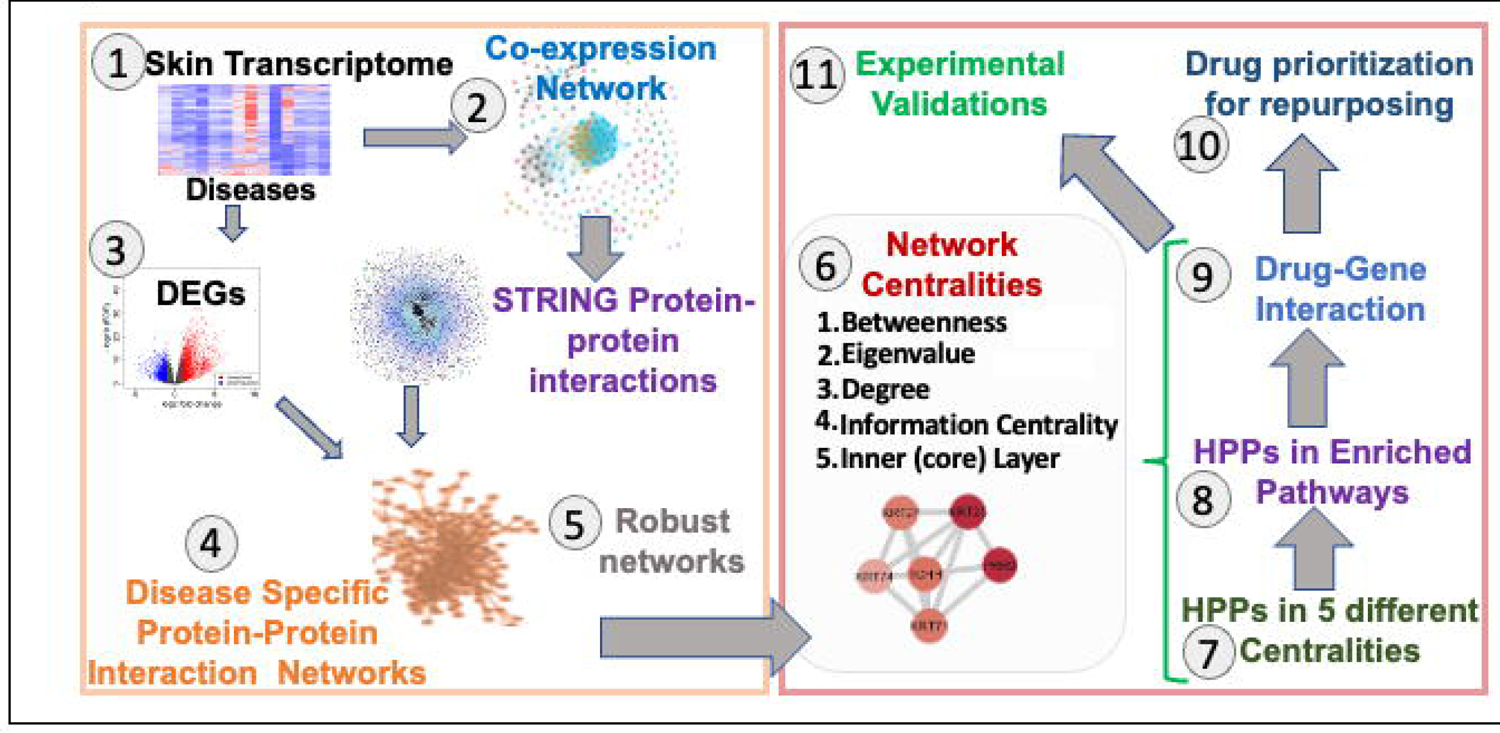
Framework to Prioritize Genes/Pathways from Transcriptome. A Schematic representation of the implemented integrative multi-omics framework utilizing the transcriptome datasets and extrapolating the network-based analysis pipeline. Eleven different steps need to be followed to identify the most appropriate drug-gene interaction for any immunogenetic-related disease.

### Different chronic inflammatory skin diseases display unique and conserved transcriptome activity

The conventional transcriptome-based analysis demonstrates the expression behavior in disease for the corresponding cohort of samples. To begin, we identified the total number of DEGs from the transcriptomics (RNA-Seq and microarray) datasets in inflammatory skin diseases including acne, AD, AK, CD, HS, ICD, PS, and three types of RS; erythematotelangiectatic rosacea (ER), phymatous rosacea (PhyR), and papulopustular rosacea (PapR) with a threshold (log2FC ≥|1|; FDR <0.05). Based on the transcriptomics platform and diseases, we captured a diverse number of DEGs per disease (Figure 2A, Table S1). For example, ICD has 65 up- and 47 down-regulated genes, whereas most DEGs were discovered in PS disease with 1763 up- and 3243 down-regulated genes, respectively. Whereas, acne, CD, and HS have a similar breakdown of DEGs (Table S1). Afterward, we performed the functional analysis for DEGs in each disease to explore the activated/inhibited pathways in those datasets. We report that the Th1 pathway, dendritic cell maturation, Th17 activation pathway, interferon signaling, Th2 pathway, IL-8 signaling, GP6 signaling pathway, CD28 signaling in T helper cells, HOTAIR regulatory pathway, NF-κB signaling are some of the significant activated canonical pathways in these chronic inflammatory skin diseases (BH *P*-value <0.05; Figure 2B, Table S1). Interestingly, we found that the senescence pathway, 1L-7 signaling, and IL-13 signaling are activated only in acne and AK, whereas B Cell receptor signaling was activated in HS, PapR, and CD.

**Figure 2.**
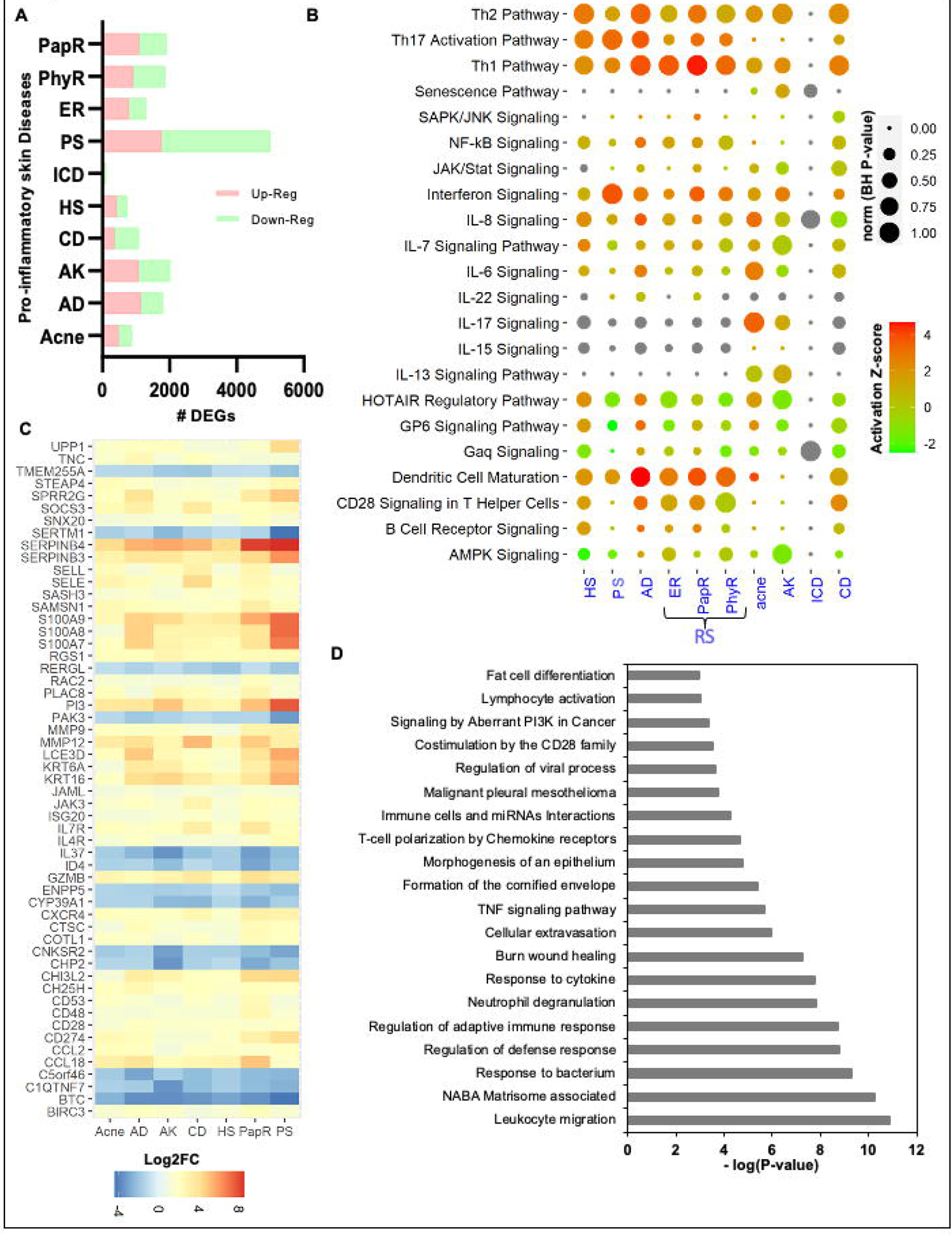
Differential expression profiles in eight inflammatory skin diseases. **A.** The total number of differentially expressed genes (DEGs) in inflammatory skin diseases including acne, atopic dermatitis (AD), actinic keratoses (AK), contact dermatitis (CD), hidradenitis suppurativa (HS), irritant contact dermatitis (ICD), psoriasis (PS), and three types of rosacea (RS; erythematotelangiectatic rosacea (ER), phymatous rosacea (PhyR), and papulopustular rosacea (PapR) with a threshold (log2FC ≥|1|; FDR <0.05). Upregulated DEGs are marked in light-red and down-regulated DEGs are marked in light green. **B.** Significantly activated and inhibited canonical pathways in different inflammatory skin diseases (BH *P*-value <0.05). **C.** Differential expression profile of total common DEGs (55) in eight inflammatory skin diseases. **D.** The enriched functional pathways and ontologies of 55 common DEGs across eight inflammatory skin diseases (P-value <0.05).

To identify the common transcriptome activity among each disease, we performed a commonality analysis. We found that 55 DEGs (including CD28, CD48, CD53, ID4, IL37, IL4R, IL7R, JAK3, KRT16, KRT6A, PI3, S100A7, S100A8, S100A9, SERPINB3, SERPINB4, and UPP1) are common in seven inflammatory skin diseases, except irritant contact dermatitis (ignored due to significantly a smaller number of DEGs reported) (Figure 2C, Table S1). These are the core transcriptome signatures across disease and involved in leukocyte migration, NABA Matrisome associated, response to the bacterium, regulation of defense response, regulation of adaptive immune response, neutrophil degranulation, cytokine response, burn wound healing, cellular extravasation, TNF signaling, formation of the cornified envelope, morphogenesis of epithelium, T-cell polarization by chemokine receptors, Immune cells and miRNAs interactions, costimulation by the CD28 family, signaling by aberrant PI3K in cancer, and lymphocyte activation (*P*-value <0.05; Figure 2D, Table S1).

These initial results suggest that there is a huge complexity among chronic inflammatory skin diseases with only a few shared and distinct genes and associated pathways. Thus, we explored the system biology-based methods to unravel intricate relationships, novel gene regulators, and master regulators of pathways altered in these diseases.

### The correlation method predicted novel genes involved in four (AD, HS, PS, and RS) chronic skin diseases

The correlation-based gene clustering is used to reduce the high-dimensional transcriptomics datasets for easier human interpretations by clustering genes into several groups based on their expression profile. GCN-based clustering is also a subtype of conventional clustering with reduced complexity through network representation. The resultant GCN can be a source point to predict the relationships among genes in a biological pathway through the guilt-by-association concept^32^. However, the main challenge is to restrict the spurious correlations, which can be achieved to a great extent by using datasets with a higher number of samples/conditions, robust GCN network construction algorithms, and standard parameters for each experiment. Thus, we performed our correlation-based analysis through WGCNA with the same parameters of power-cutoff, weight-cutoff, and clustering algorithm for eight inflammatory skin disease transcriptomes^27^. There are eight resultant GCNs of which acne (320 nodes and 264 edges) is the smallest, HS is the largest by an edge (4,162 nodes and 420,422 edges) and, PS (5,465 nodes and 132,849 edges) is the largest by nodes. AK, CD, and ICD have a comparable number of nodes and an edge. (Figure 3A, Table S2).

**Figure 3.**
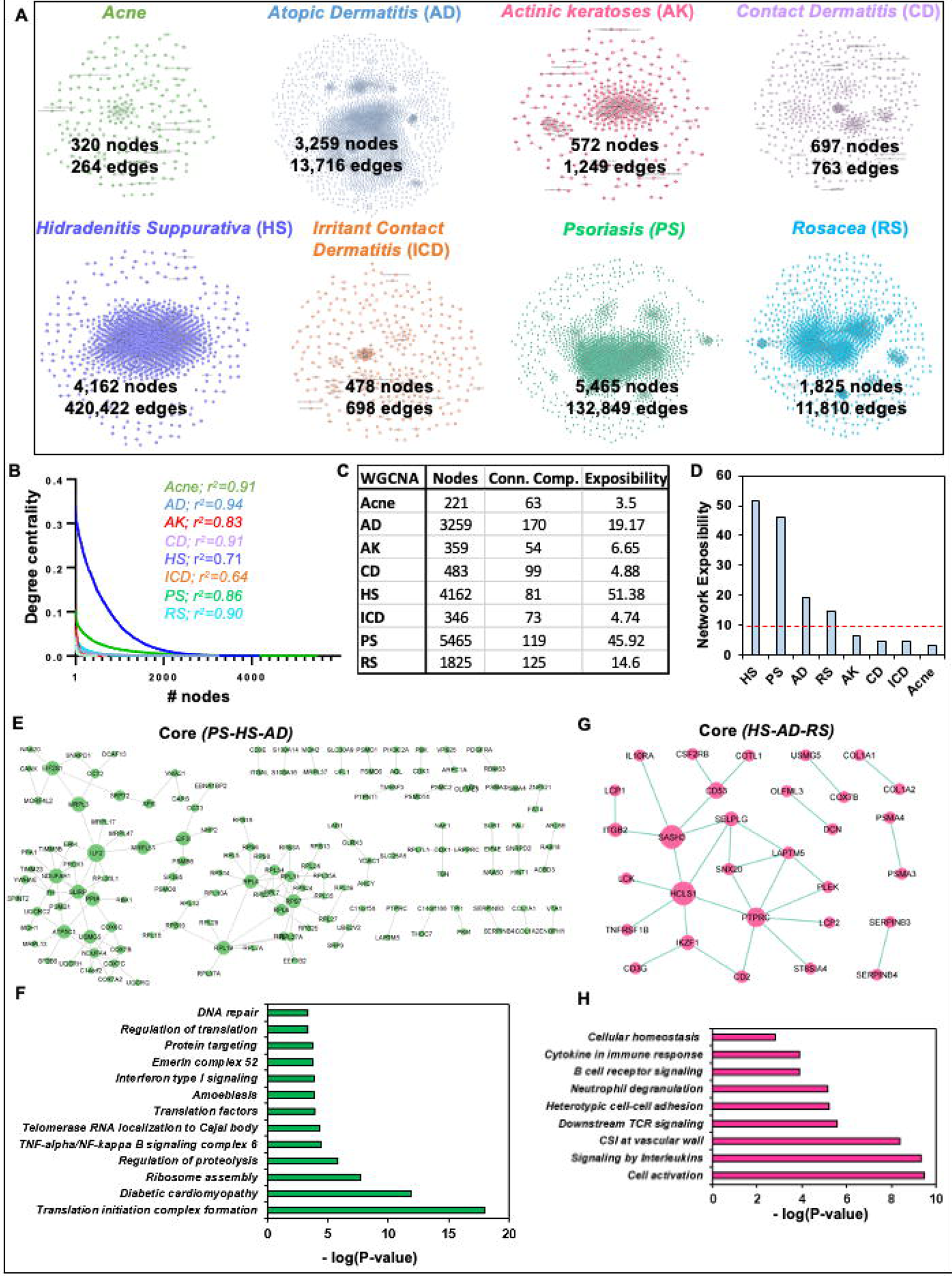
The co-expression networks highlight core genes in several inflammatory skin diseases. **A.** Different co-expression networks were constructed with the same parameters through WGCNA for eight inflammatory skin diseases with the corresponding number of nodes and edges. The networks are acne (320 nodes and 264 edges), atopic dermatitis (AD; 3,259 nodes and 13,716 edges), actinic keratoses (AK; 572 nodes and 1,249 edges), contact dermatitis (CD; 697 nodes and 763 edges), hidradenitis suppurativa (HS; 4,162 nodes and 420,422 edges), irritant contact dermatitis (ICD; 478 nodes and 698 edges), psoriasis (PS; 5,465 nodes and 132,849 edges), and rosacea (RS; 1,825 nodes and 11,810 edges). **B.** The power-law distribution of eight co-expression networks to measure their scale-freeness. **C.** The table illustrates the total number of nodes and connected components to determine the network exposibility for each co-expression network. **D.** The network exposibility distribution for eight co-expression networks to illustrate the robust networks to study further with parameter (network exposibility >10). **E.** The core co-expression network among psoriasis, hidradenitis suppurativa, and atopic dermatitis. **F.** The gene ontology analysis of enriched pathway of core co-expression network genes among PS, HS, and AD (*P*-value <0.05). **G.** The core co-expression network among HS, AD, and RS. **H.** The gene ontology analysis of enriched pathway of core co-expression network genes among hidradenitis suppurativa, atopic dermatitis, and rosacea (P-value <0.05).

To begin our GCNs analysis, we performed several network centrality analyses including the degree to check the scale-free properties of each network^33^. As expected, we found that all eight GCNs; Acne, AD, AK, CD, HS, ICD, PS, and RS follow scale-free properties calculated by the power-law distribution with *r*^2^ values 0.94 for AD and 0.64 for ICD (Figure 3B, Table S2). Though most of these GCNs are biologically relevant through power-law distribution, they don’t seem comparable to each other. To check the extent of coverage in a network, we calculated the network exposibility as described in the methods section. We found that four GCNs corresponding to acne, AK, CD, and ICD have a small number of nodes and a high number of connected components, whereas the remaining four other GCNs related to AD, HS, PS, and RS have a high number of nodes but a smaller number of connected components comparably (Figure 3C, Table S2). To determine the exposibility-based reliability of eight GCNs, we set a network exposibility threshold ≥ 10 and filtered our four disease-specific GCNs including HS, PS, AD, and RS for further analysis (Figure 3D).

It is well reported that some of the immune-mediated diseases including psoriasis, HS, and AD share most of their disease-associated pathways and comorbidities with each other^34,35^. Therefore, to understand the shared gene patterns among these diseases, we explored the core co-expression network of four diseases. Interestingly, we report a core co-expression network among PS, HS, and AD with 145 genes and 161 associations (Figure 3E, Table S2). Interestingly, some of these genes are RPS-, RPL-, MRPL-, EIF-, COX-family genes, ILF2, PPIA, NDUFAB1, NDUFA4, ATP5C1, and FH.

Furthermore, to verify the biological associations of 145 core genes, we performed gene ontology analysis. The ontology analysis identified the enriched pathways including translation initiation complex formation, diabetic cardiomyopathy, ribosome assembly, regulation of proteolysis, TNF-alpha/NF-kappa B signaling complex 6, telomerase RNA localization to Cajal body, translation factors, amoebiasis, Interferon type I signaling, Emerin complex-52, protein targeting, and DNA repair (*P*-value <0.05; Figure 3F, Table S2). Similarly, we explored the core co-expression network among PS, HS, and RS. In this analysis, we found a small core network encompassing 30 genes and 30 associations (Figure 3G, Table S2). Interestingly, some of these genes are IKAROS Family Zinc Finger 1 (IKZF1), CD2 Molecule (CD2), CD53 Molecule (CD53), SAM And SH3 Domain Containing 3 (SASH3), Lysosomal Protein Transmembrane 5 (LAPTM5), Interleukin 10 Receptor Subunit Alpha (IL10RA), Protein Tyrosine Phosphatase Receptor Type C (PTPRC), and Pleckstrin (PLEK). Furthermore, to verify the biological associations of 30 core genes in HS, AD, and RS, we performed gene ontology analysis. We found that most of the enriched pathways are cell activation, signaling by Interleukins, CSI at the vascular wall, downstream TCR signaling, heterotypic cell-cell adhesion, neutrophil degranulation, B cell receptor signaling, cytokine in immune response, and cellular homeostasis (*P*-value <0.05; Figure 3H, Table S2). These results suggest the significance of correlation-based GCNs construction and analyses to identify the emerging and shared players in associated pathways among inflammatory skin diseases.

### Interactome analysis identified the central proteins in psoriasis, hidradenitis suppurativa, atopic dermatitis, and rosacea

The molecular mechanisms and intricacies of biological processes associated with immune-mediated skin diseases are mostly determined by protein interactions in certain pathways^36^. Given that the coverage of DEGs from transcriptome to differentially expressed proteins is a maximum of only ∼60%, we used the disease-specific co-expressed gene list to extract the PPI network. These interactions can be more relevant in any stress condition if they are also part of corresponding GCNs by satisfying the guilt-by-association assumption^37^. To explore these possibilities, we constricted the disease-specific PPI network by integrating the co-expressed genes into the known interactions from the STRING database^29^. The resultant PPI networks/interactomes for these four diseases include PS with the greatest number of nodes (3,622 nodes and 33,173 edges) and RS with the least number of nodes (851 nodes and 3,234 edges) (Figure 4A, Table S3). It is reported that some of the molecular pathways and disease comorbidities are shared among psoriasis, hidradenitis suppurativa, atopic dermatitis, and rosacea^1^. To identify the core proteins in four PPIs, we performed the commonality analysis and found that 53 proteins are shared among these four inflammatory skin diseases (Figure 4B, Table S3). Some of these proteins are CD2, CD3D, CD53, CD96, CDK1, COL1A1, COL1A2, COL3A1, COX5A, COX7B, CXCR4, DCN, FH, IKZF3, IL10RA, ITGAL, KRT6A, KRT6B, LAPTM5, and LCK, which participate in lymphocyte activation, diabetic cardiomyopathy, neutrophil degranulation, membrane raft distribution, lymphocyte-mediated immunity, TYROBP causal network in microglia, response to molecule of bacterial origin, microglia pathogen phagocytosis pathway, homeostasis of several cells, formation of the cornified envelope, negative regulation of defense response, cell cycle phase transition, positive regulation of endopeptidase activity, regulation of glial cell differentiation, acute viral myocarditis, human cytomegalovirus infection, and immune effector process (Figure S1, Table S3).

**Figure 4.**
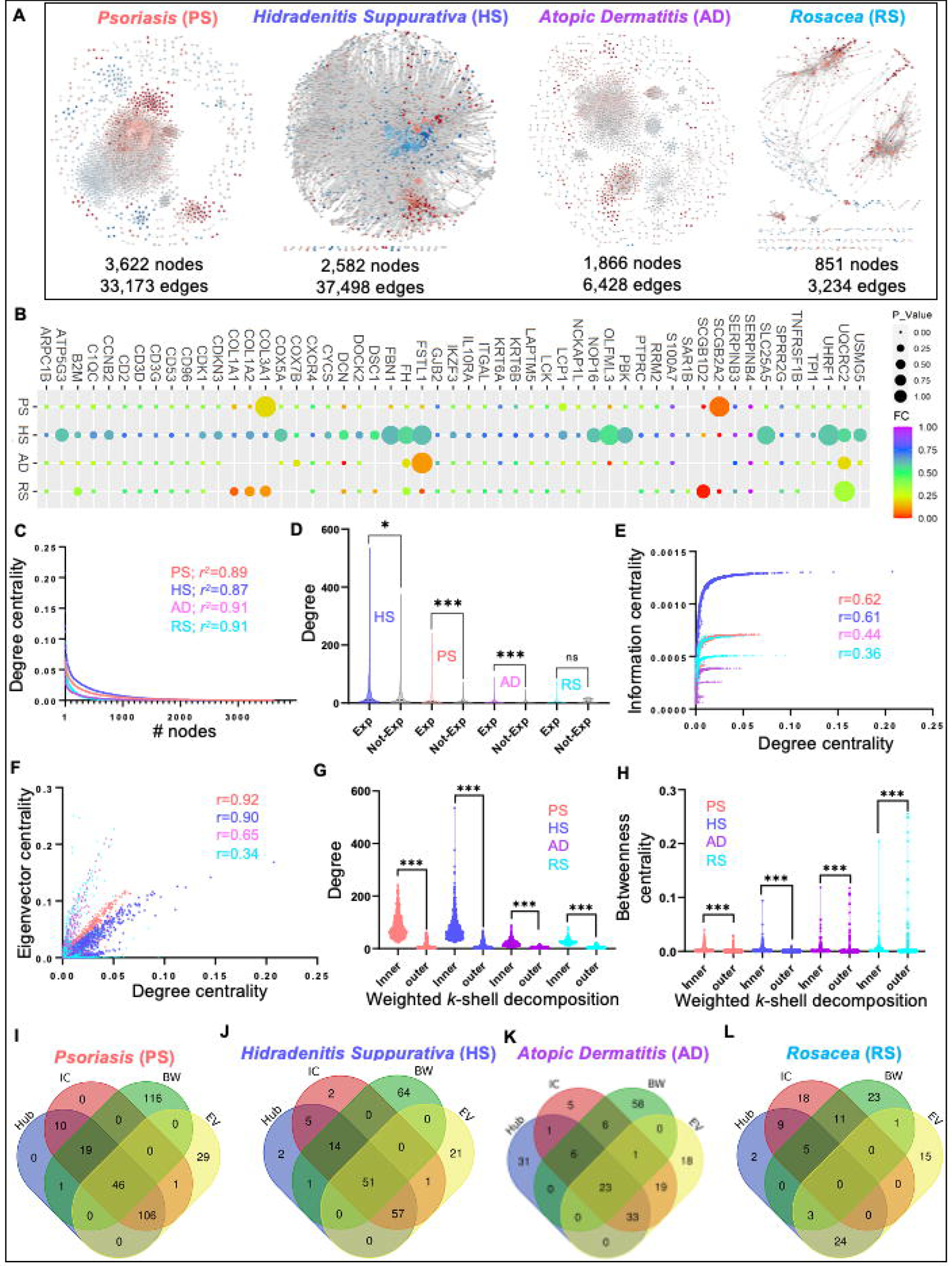
The protein-protein interaction (PPI) networks describe the structural topology of proteins in four inflammatory skin diseases. **A.** Different PPI networks were constructed with a compendium of human interactome extracted by co-expression network nodes for four inflammatory skin diseases with a corresponding number of nodes and edges. The PPI networks are psoriasis (PS; 3,622 nodes and 33,173 edges), hidradenitis suppurativa (HS; 2,582 nodes and 37,498 edges), atopic dermatitis (AD; 1,866 nodes and 6,428 edges), and rosacea (RS; 851 nodes and 3,234 edges). **B.** The shared proteins in all four PPI and their expression pattern across four diseases. **C.** The power-law distribution calculation of eight co-expression networks to measure their scale-freeness. **D.** The degree distribution of expressed genes (Exp) and not expressed genes (Not-Exp) in four disease-specific PPIs. **E-F.** The correlation plots illustrate the distribution among degree, information, and eigenvector of all four PPI networks. **G-H.** The distribution of inner and outer layer proteins with a degree as well as betweenness centrality in all four PPI networks. **I-L.** The shared and unique proteins with significantly high centralities (top 5%) for individual disease-specific PPI networks.

The disease-specific interactomes position their important proteins in a particular arrangement to communicate throughout the network most efficiently. To explore these structural arrangements, we performed network analysis on four disease-specific PPIs including PS, HS, AD, and RS. To begin our analysis first, we tested the biological relevance of interactomes by performing the power-law distribution analysis of four PPIs to verify their scale-freeness^33,38^. Interestingly, we report that all four PPIs for PS, HS, AD, and RS follow scale-free properties calculated by the power-law distribution and the *r*^2^ values are similar (Figure 4C, Table S3). Previously it has been reported that the network-based method has successfully highlighted the central and core proteins associated with disease pathogenesis^9,36,38^. Therefore, we leveraged a part of this analysis in our framework and analyzed four PPIs with a degree, betweenness centrality, eigenvector, information centrality, and weighted *k*-sell decomposition method. First, we explored the degree distribution of expressed genes (Exp; *FDR*<0.05) and non-expressed (Not-Exp; *FDR*>0.05) for each disease. Interestingly, we found that expressed genes of all four diseases namely HS, PS, AD, and RS encompass a high degree distribution than their non-expressed counterparts (Figure 4D, Table S3), Mann-Whitney test *P*-value (HS<0.05, PS< 0.0001, AD<0.0001, and RS>0.66).

Previously, we have demonstrated that some of the highly connected genes also possess other network properties, which make them extremely vulnerable during disease pathogenesis^9,18^. Therefore, we calculated the correlation between degree centrality and other centralities including information, eigenvector, and betweenness centralities of all four PPI networks. Interestingly, we found that the degree of PS and HS are strongly correlated with information centrality with *r*>0.6 for both, whereas this correlation is not strong in AD and RS with *r*<0.5 (Figure 4E). Similarly, we computed the correlation among degree and eigenvector centrality and reported that PS, HS, and AD have the strongest correlation (*r*<0.5), whereas RS has the weakest correlation (*r*<0.5, Figure 4F). Differently, the correlation between degree and betweenness is very poor in PS, AD, and RS (*r*<0.5) and strong in HS (*r*>0.5) (Figure S1B). Further, we decomposed the PPI networks into *k*-shells through the Weighted k-shell decomposition method to identify the hidden genes, which the conventional centralities fail to identify^13^. Next, we identified the inner layer and peripheral (outer) layer proteins in each PPI as mentioned by Ahmed et al., 2018^13^. We report that there are 461, 573, 393, 95 proteins in the inner layer and 3135, 1982, 1466, and 754 proteins in the outer layer of PS, HS, AD, and RS, respectively (Table S3). Moving forward, we explored the degree and betweenness centrality distributions of the inner layer and outer layer proteins in four PPIs. We report that inner-layer proteins possess a significantly higher degree compared to outer-layer proteins (Figure 4G; Mann-Whitney test *P*-value <0.0001, Table S3). However, we do not see much difference in the betweenness centrality distribution of inner and outer layer proteins (Figure 4H; Mann-Whitney test P-value <0.0001). These analyses confirm that some of the proteins of PPIs share different types of high centrality values, which can be central in the disease pathogenesis. Additionally, we investigated the high centrality proteins (significant proteins) by selecting the top5% of high centralities i.e Hub (degree), bottleneck (Betweenness, BW), information centrality (IC), and eigenvector centrality (EV) for individual disease-specific PPI network. We demonstrate that 46, 51, and 23 proteins are shared by all four centralities in PS, HS, and AD, respectively (Figure 4I-K, Table S3). Whereas there was not a single protein shared by all four centralities in RS (Figure 4L, Table S3). Interestingly, we also found that most of the high betweenness centrality (bottleneck) proteins are not shared by any other centrality. These analyses indicate that most of these bottlenecks are part of smaller subnetworks in four PPIs.

### Network centrality-based prioritization of proteins and pathways in chronic inflammatory skin diseases

It has been previously described in several instances that in the disease-specific interactomes, the most significant contributing proteins have a high degree (connections) and high betweenness (bottlenecks) against other proteins throughout the PPI network^14–16,36,39^. Further, the analysis was expanded to other centralities including information and eigenvector centralities, which improved the identification of significantly contributing proteins in the interactome^9^. Inspired by these studies, we identified the significant proteins for disease-specific PPI networks of PS. HS, AD, and RS individually. The top 5% of centrality value nodes with the degree, betweenness, eigenvector, information centrality, and the inner layer proteins from the weighted k-shell decomposition method were identified as significant proteins. Next, we put our effort into classifying these significant proteins as regulators in four skin diseases in this study and termed them as HPPs or significant regulators (Figure 5A, Table S4). As a result, we identified 55 HPPs that contribute significantly to the pathogenesis of four selected diseases (Table S4). Further, we explored the regulator activity of these 55 HPPs and found that most of these proteins are significantly activated in at least seven inflammatory skin diseases (Figure 5B, BH *P*-value <0.05; Activation z-score > |1|; Table S4). Interestingly, some of these 55 HPPs are involved in RTKs Signaling, proteoglycans in cancer, cell morphogenesis, hemopoiesis, PID CXCR4 pathway, epithelial cell differentiation, Cytokine Signaling in the Immune system, adherens junction, regulation of kinase activity, cellular response to lipid, head, and neck SCC, response to wounding, Hippo signaling regulation, Interferon type I signaling, PI3K-Akt signaling, leukocyte activation, response to estradiol, lymphocyte activation, Th1, and Th2 cell differentiation, and NK cell-mediated cytotoxicity (Figure 5C, Table S4; *P*-value <0.05). Most of these HPPs and associated pathways are known for their major contribution to several immune-mediated diseases^5–7,20,25,36,38,40,41^. Moving forward, we investigated the reported potential therapeutics for candidate 55 HPPs, which will help in designing highly effective drug repurposing strategies. We extracted the Drug-Gene interaction network pairs from DGIdb^30^ for 55 HPPs with available literature publications. We found a total of 32 HPPs have a known compendium of 199 drug compounds with 237 interactions (Figure 5D, Table S4). Interestingly, we report that some 55 HPPs i.e., CD2, LCK, STAT1, TNFRSF1B, IKZF1, APP, and BMP7 can be a high-priority drug target for several of these chronic skin diseases.

**Figure 5.**
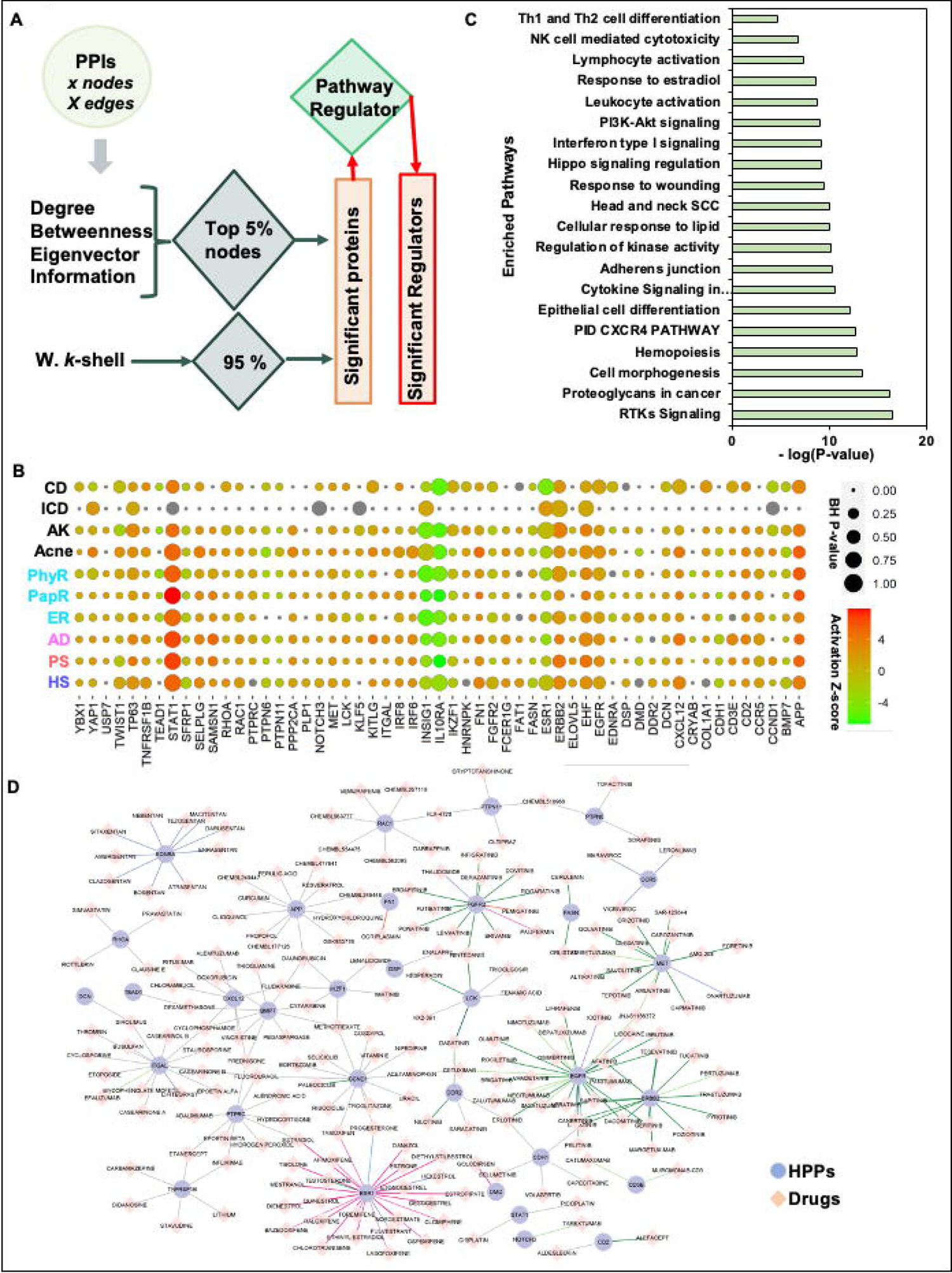
The identification of high-priority proteins (HPPs) from disease-specific protein-protein interaction (PPI) networks and their regulator activity. **A.** The framework to identify the HPPs for each PPI network. The top 5% of centrality value nodes with a degree, betweenness, eigenvector, information centrality, and the inner layer proteins from the weighted k-shell decomposition method were identified as significant proteins. If these significant proteins are also significantly activated or inhibited regulators in each of the eight inflammatory skin diseases are termed HPPs or significant regulators. **B.** Our network-centric approach identified 55 HPPs and their activity across inflammatory skin diseases. **C.** The gene ontology analysis of the enriched pathway of 55 HPPs (P-value <0.05). **D.** Drug-gene interaction network pairs for HPPs and chemical compounds with publication listed in DGIdb.

### IKZF1 is a potential therapeutic target for several inflammatory skin diseases

Given the overlap in the molecular profiles of multiple inflammatory skin diseases, we explored the shared significant regulator proteins in HS, PS, AD, and RS. For this, we complied all the significant proteins based on five centrality methods (degree, betweenness, information, eigenvector centrality, and weighted *k*-shell decomposition) for each of these diseases. As a result, we have identified a list of 236, 195, 168, and 72 significant proteins respectively in PS, HS, AD, and RS (Figure 6A, Table S4). Afterward, we studied the distribution of shared and unique significant proteins in each PPI network and found that six proteins (NDUFAB1, MRPL3, DDX1, EIF2S1, SLIRP, and ATP5C1) are contributing significantly to these four diseases. Whereas there is only one protein IKZF1 contributing significantly to HS, AD, and RS with high network centrality values, we explored the drug-gene interaction of IKZF1. In this regard, six drugs including lenalidomide, daunorubicin, methotrexate, cytarabine, imatinib, and fludarabine are known to target IKZF1 (Figure 6B). Interestingly, we also found that daunorubicin also targets APP and BMP7 which are also part of predicted 55 HPPs. Similarly, methotrexate also targets CCND1 and BMP7, and cytarabine targets BMP7 too (Figure 5D). Thereafter, we extracted all the possible interactions of IKZF1 from HS-PPI, HS-GCN, and known TF-target relationships to explore the extrapolating effect of a drug on the IKZF1 and its partners. We found that IKZF1 can be a really good candidate as it acts as a master regulator by interacting with 19 targets, 62 proteins, and 88 co-expressed genes (Figure 6C, Table S4). We hypothesized that the activities of these proteins and co-expressed genes can be altered by targeting IKZF1 with a specific drug, such as lenalidomide. As IKZF1 is shared among three of these skin diseases, we surveyed the expression pattern of IKZF1 interacting and co-expressing genes in HS, PS, AD, and RS. Interestingly, we found that most of the IKZF1 interacting partners; 19 targets, 62 proteins, and 88 co-expressed genes follow a similar expression pattern across four inflammatory skin diseases (Figure 6 D-F, Table S4).

**Figure 6.**
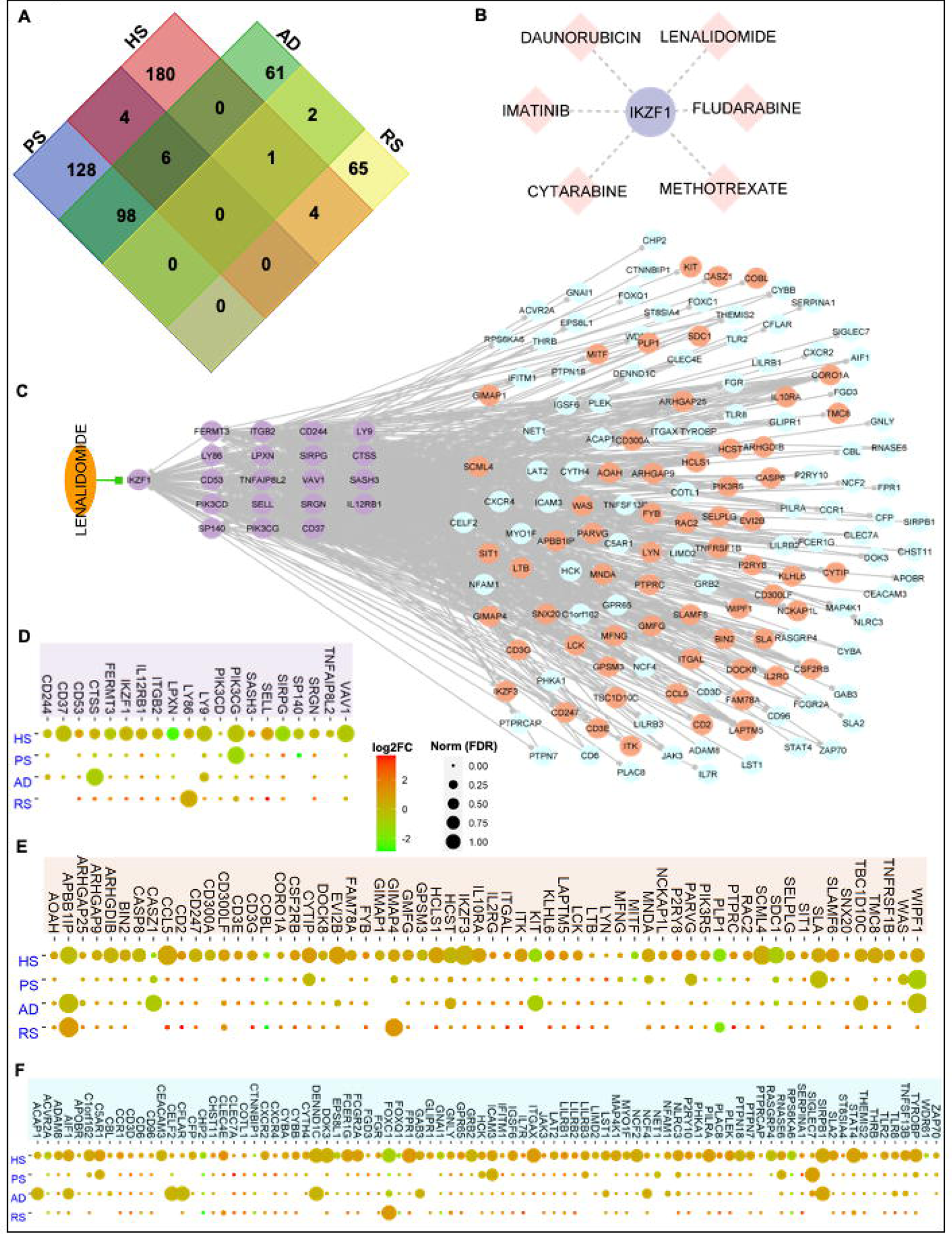
The identification of shared significant regulator proteins in four inflammatory skin diseases. **A.** The Venn diagram represents the distribution of shared and unique significant proteins in each PPI network. Interestingly, IKZF1 protein is shared among hidradenitis suppurativa (HS), atopic dermatitis (AD), and rosacea (RS) with high network centrality values. **B.** The six drugs are known to target IKZF1. **C.** The IKZF1 acts as a master regulator by interacting with 19 targets, 62 proteins, and 88 co-expressed genes. Activities of these proteins can be altered by targeting IKZF1 with the drug Lenalidomide. D-F. The expression profile of IKZF1 interacting partners; 19 targets, 62 proteins, and 88 co-expressed genes in four inflammatory skin diseases including HS, PS, AD, and RS.

Taken together, our study identified high-value proteins involved in the pathogenesis of chronic skin diseases including HS, PS, AD, and RS. Additionally, we designed a robust framework to identify significant contributors utilizing transcriptome datasets and integrative multi-omics approaches supported by network systems biology for more accurate predictions of regulators in most chronic skin diseases.

## DISCUSSION

Almost 1/5^th^ of the human population across the world is affected by some type of non-communicable chronic inflammatory disease. The disease manifestation and immune regulatory signatures are shared among some of these inflammatory skin diseases. The high-throughput ‘omics technology and robust multi-omics network integration techniques showed immense potential to accelerate more comprehensive and system-wide discoveries in the pathogenesis of complex inflammatory skin diseases. Here, our designed robust framework identified significant contributors from different transcriptome datasets and integrative multi-omics supported the network systems biology for more accurate predictions of significant regulators of these inflammatory skin diseases (namely: acne, atopic dermatitis (AD), actinic keratoses (AK), contact dermatitis (CD), irritant contact dermatitis (ICD), psoriasis (PS), hidradenitis suppurativa (HS), and three types of rosacea (RS)). Our conventional and unconventional biological data analysis approach extracted the shared and unique biomolecular (gene/protein/regulator) signatures and biological processes in four (PS, PS, AD, and RS) of these eight inflammatory skin diseases. Further, systems genetics and network biology-driven analyses were instrumental in prioritizing the most relevant proteins (55 high-priority proteins; HPPs) across four of these inflammatory skin diseases. These HPPs can serve as a template to study the impact and response of disease pathogenesis of these inflammatory skin diseases, and putative drug targets strategies either as standalone or combinatorial approaches. Overall, this study employs advanced and powerful network systems-based analyses that combine integrative multi-omics to establish a mutual understanding of inflammatory mechanisms in four skin diseases including PS, PS, AD, and RS.

In the last two-decade, PS has been one of the most studied inflammatory skin diseases, which shares a significant number of immune-mediated pathways with other inflammatory diseases including acne, AD, AK, HS, and RS. For example, similar to psoriasis, targeting Th2 cells, IL-4, and IL-13 have been the most efficacious (50-70%), and advanced therapy for AD^40^ and Janus kinase (JAK) inhibitors has been demonstrated to suppress the cytokine responses more effectively and are also a powerful treatment strategy for AD^40^. HS is also an inflammatory skin disease implicated by the pathogenesis of neutrophilic inflammation, dysbiosis, TNF, interferon responses, hair- and skin-gland abnormalities, autoantibodies, and plasma cells^41^. Our multi-omics analysis identified the shared molecules, regulators, and pathways at transcriptome and interactome layers in four diseases including TNF-alpha/NF-kappa B signaling, translation factors, amoebiasis, Interferon type I signaling, Cytokine signaling, and interleukin signaling. These pathways have been under investigation to manage/treat several immune system diseases.

Our network analytics utilizing different conventional and improved centralities appreciably identified the central proteins of these skin disease-specific interactomes as well as substantially improved the classification of significant proteins (55 HPPs) in the pathogenesis of PS, HS, AD, and RS. The functional analysis of these HPPs further highlighted the Cytokine Signaling, adherens junction, response to wounding, Hippo signaling regulation, Interferon type I signaling, PI3K-Akt signaling, leukocyte activation, Th1, and Th2 cell differentiation, NK cell-mediated cytotoxicity to name a few. Many of these pathways have been validated in various earlier investigations^36,40,42^. These findings demonstrate the effectiveness of a network centrality-based framework to untangle the underlying players of these diseases.

Our integrative multi-omics framework has identified IKZF1 as a significant shared regulator among HS, AD, and RS. Remarkably, IKZF1 is associated with chromatin remodeling and lymphocyte differentiation which can influence the disease pathogenesis. A considerable amount of IKZF1 interacting partners in HS interactome and co-expression networks suggest the involvement of this transcription factor in the immune environment modulation in the disease pathogenesis. Interestingly, we also found most immune-system pathways including cellular response to cytokine stimulus, leukocyte activation, neutrophil degranulation, inflammatory response, TCR pathway, CXCR4 pathway, hemostasis, and microglia pathogen phagocytosis pathway is enriched among IKZF1 interactors. The specific role of neutrophils, macrophages, B cells, and plasma cells has been identified in HS pathogenesis^42–44^. Similarly, CXCR4 pathways is considered important in regulating B cell functions, homing of plasma cells, and in the migration of T cells^45,46^. Interestingly, many small molecule drugs are known modulators of IKZF1 response. It is likely that these or similar molecules may find a place in developing therapeutic innervations of these complex inflammatory skin diseases. Overall, our integrative network-based multi-omics framework identified the underlying regulators, and pathways common among these diverse proinflammatory diseases.

This study also has limitations as available datasets for bulk transcriptome-driven/disease integrative multi-omics inflammatory skin diseases, will not generate the most reliable predictions of significant regulators, yet this framework as suggested here can be applied for multiple transcriptome datasets of a single disease for more accurate predictions. Furthermore, expanding the current version of transcriptome-driven integrative systeomics to the next level by implementing other multi-layered datasets including systems genomics, epigenomics and the emergence of new technologies like single-cell RNA sequencing, and spatial transcriptomics will uncover the high-resolution molecular pathogenesis, pathways, and cell-cell communications in chronic inflammatory skin disease.

---

**Figure S1.** The significant proteins in four inflammatory skin diseases. **A.** The gene ontology of 53 shared proteins in four disease analyses identified lymphocyte activation, diabetic cardiomyopathy, neutrophil degranulation, membrane raft distribution, lymphocyte-mediated immunity, TYROBP causal network in microglia, response to molecule of bacterial origin, microglia pathogen phagocytosis pathway, homeostasis of several cells, formation of the cornified envelope, negative regulation of defense response, cell cycle phase transition, positive regulation of endopeptidase activity, regulation of glial cell differentiation, acute viral myocarditis, human cytomegalovirus infection, and immune effector process as significantly enriched pathways (*P*-value<0.05). **B.** The correlation plot illustrates the distribution among degree and betweenness centrality of all four PPI networks.

## DECLARATIONS

### Authors’ contributions

M.S.M., B.M., and M.A. conceived the project. B.M. performed all the computational, network-based, and statistical analyses. B.M. wrote the first draft and final version of the manuscript. All the authors discussed the results, critically reviewed the manuscript, and provided valuable comments/edits.

### Availability of data and materials

All datasets used and generated from this study are accessible through Table S files. This study did not generate new unique reagents. Requests for materials and communications with the journal should be addressed to M.S.M. and M.A.

## References

1 Wang, L., Wang, F. S. & Gershwin, M. E. Human autoimmune diseases: a comprehensive update. J Intern Med 278, 369–395 (2015). 10.1111/joim.12395

2 Schwingen, J., Kaplan, M. & Kurschus, F. C. Review-Current Concepts in Inflammatory Skin Diseases Evolved by Transcriptome Analysis: In-Depth Analysis of Atopic Dermatitis and Psoriasis. Int J Mol Sci 21 (2020). 10.3390/ijms21030699

3 Carretero, M. et al. Differential Features between Chronic Skin Inflammatory Diseases Revealed in Skin-Humanized Psoriasis and Atopic Dermatitis Mouse Models. J Invest Dermatol 136, 136–145 (2016). 10.1038/JID.2015.362

4 Boehncke, W. H. & Schon, M. P. Psoriasis. Lancet 386, 983–994 (2015). 10.1016/S0140-6736(14)61909-7

5 Armstrong, A. W. & Read, C. Pathophysiology, Clinical Presentation, and Treatment of Psoriasis: A Review. JAMA 323, 1945–1960 (2020). 10.1001/jama.2020.4006

6 Goldburg, S. R., Strober, B. E. & Payette, M. J. Hidradenitis suppurativa: Current and emerging treatments. J Am Acad Dermatol 82, 1061–1082 (2020). 10.1016/j.jaad.2019.08.089

7 Kim, J. & Krueger, J. G. Highly Effective New Treatments for Psoriasis Target the IL-23/Type 17 T Cell Autoimmune Axis. Annu Rev Med 68, 255–269 (2017). 10.1146/annurev-med-042915-103905

8 Weidinger, S., Beck, L. A., Bieber, T., Kabashima, K. & Irvine, A. D. Atopic dermatitis. Nat Rev Dis Primers 4, 1 (2018). 10.1038/s41572-018-0001-z

9 Kumar, N., Mishra, B., Mehmood, A., Mohammad, A. & Mukhtar, M. S. Integrative Network Biology Framework Elucidates Molecular Mechanisms of SARS-CoV-2 Pathogenesis. iScience 23, 101526 (2020). 10.1016/j.isci.2020.101526

10 von Mering, C. et al. Comparative assessment of large-scale data sets of protein-protein interactions. Nature 417, 399–403 (2002). 10.1038/nature750

11 Alm, E. & Arkin, A. P. Biological networks. Curr Opin Struct Biol 13, 193–202 (2003). 10.1016/s0959-440x(03)00031-9

12 Vidal, M., Cusick, M. E. & Barabasi, A. L. Interactome networks and human disease. Cell 144, 986–998 (2011). 10.1016/j.cell.2011.02.016

13 Ahmed, H. et al. Network biology discovers pathogen contact points in host protein-protein interactomes. Nat Commun 9, 2312 (2018). 10.1038/s41467-018-04632-8

14 Dai, H., Zhou, J. & Zhu, B. Gene co-expression network analysis identifies the hub genes associated with immune functions for nocturnal hemodialysis in patients with end-stage renal disease. Medicine (Baltimore) 97, e12018 (2018). 10.1097/MD.0000000000012018

15 Liu, J. et al. Identification of hub genes and pathways associated with hepatocellular carcinoma based on network strategy. Exp Ther Med 12, 2109–2119 (2016). 10.3892/etm.2016.3599

16 Liu, J., Jing, L. & Tu, X. Weighted gene co-expression network analysis identifies specific modules and hub genes related to coronary artery disease. BMC Cardiovasc Disord 16, 54 (2016). 10.1186/s12872-016-0217-3

17 Mishra, B., Athar, M. & Mukhtar, M. S. Transcriptional circuitry atlas of genetic diverse unstimulated murine and human macrophages define disparity in population-wide innate immunity. Sci Rep 11, 7373 (2021). 10.1038/s41598-021-86742-w

18 Mishra, B., Sun, Y., Howton, T. C., Kumar, N. & Mukhtar, M. S. Dynamic modeling of transcriptional gene regulatory network uncovers distinct pathways during the onset of Arabidopsis leaf senescence. NPJ Syst Biol Appl 4, 35 (2018). 10.1038/s41540-018-0071-2

19 Blok, J. L., Li, K., Brodmerkel, C., Jonkman, M. F. & Horvath, B. Gene expression profiling of skin and blood in hidradenitis suppurativa. Br J Dermatol 174, 1392–1394 (2016). 10.1111/bjd.14371

20 Buhl, T. et al. Molecular and Morphological Characterization of Inflammatory Infiltrate in Rosacea Reveals Activation of Th1/Th17 Pathways. J Invest Dermatol 135, 2198–2208 (2015). 10.1038/jid.2015.141

21 Tsoi, L. C. et al. Atopic Dermatitis Is an IL-13-Dominant Disease with Greater Molecular Heterogeneity Compared to Psoriasis. J Invest Dermatol 139, 1480–1489 (2019). 10.1016/j.jid.2018.12.018

22 Pedersen, M. B., Skov, L., Menne, T., Johansen, J. D. & Olsen, J. Gene expression time course in the human skin during elicitation of allergic contact dermatitis. J Invest Dermatol 127, 2585–2595 (2007). 10.1038/sj.jid.5700902

23 Joly, F. et al. Photodynamic therapy corrects abnormal cancer-associated gene expression observed in actinic keratosis lesions and induces a remodeling effect in photodamaged skin. J Dermatol Sci (2018). 10.1016/j.jdermsci.2018.05.002

24 Clemmensen, A. et al. Genome-wide expression analysis of human in vivo irritated epidermis: differential profiles induced by sodium lauryl sulfate and nonanoic acid. J Invest Dermatol 130, 2201–2210 (2010). 10.1038/jid.2010.102

25 Kelhala, H. L. et al. IL-17/Th17 pathway is activated in acne lesions. PLoS One 9, e105238 (2014). 10.1371/journal.pone.0105238

26 Love, M. I., Huber, W. & Anders, S. Moderated estimation of fold change and dispersion for RNA-seq data with DESeq2. Genome Biol 15, 550 (2014). 10.1186/s13059-014-0550-8

27 Langfelder, P. & Horvath, S. WGCNA: an R package for weighted correlation network analysis. BMC Bioinformatics 9, 559 (2008). 10.1186/1471-2105-9-559

28 Shannon, P. et al. Cytoscape: a software environment for integrated models of biomolecular interaction networks. Genome Res 13, 2498–2504 (2003). 10.1101/gr.1239303

29 Szklarczyk, D. et al. STRING v10: protein-protein interaction networks, integrated over the tree of life. Nucleic Acids Res 43, D447–452 (2015). 10.1093/nar/gku1003

30 Cotto, K. C. et al. DGIdb 3.0: a redesign and expansion of the drug-gene interaction database. Nucleic Acids Res 46, D1068–D1073 (2018). 10.1093/nar/gkx1143

31 Zhou, Y. et al. Metascape provides a biologist-oriented resource for the analysis of systems-level datasets. Nat Commun 10, 1523 (2019). 10.1038/s41467-019-09234-6

32 Wolfe, C. J., Kohane, I. S. & Butte, A. J. Systematic survey reveals general applicability of “guilt-by-association” within gene coexpression networks. BMC Bioinformatics 6, 227 (2005). 10.1186/1471-2105-6-227

33 Broido, A. D. & Clauset, A. Scale-free networks are rare. Nat Commun 10, 1017 (2019). 10.1038/s41467-019-08746-5

34. Comorbidity screening in Hidradenitis Suppurativa: evidence-based recommendations from the US and Canadian Hidradenitis Suppurativa Foundations. Journal of the American Academy of Dermatology (2021).

35. New treatments in atopic dermatitis. Annals of Allergy, Asthma & Immunology 126(1) (2021).

36 Su, W., Wei, Y., Huang, B. & Ji, J. Identification of Hub Genes and Immune Infiltration in Psoriasis by Bioinformatics Method. Front Genet 12, 606065 (2021). 10.3389/fgene.2021.606065

37 van Dam, S., Vosa, U., van der Graaf, A., Franke, L. & de Magalhaes, J. P. Gene co-expression analysis for functional classification and gene-disease predictions. Brief Bioinform 19, 575–592 (2018). 10.1093/bib/bbw139

38 Manczinger, M. & Kemeny, L. Novel factors in the pathogenesis of psoriasis and potential drug candidates are found with systems biology approach. PLoS One 8, e80751 (2013). 10.1371/journal.pone.0080751

39 Wee, J. J. & Kumar, S. Prediction of hub genes of Alzheimer’s disease using a protein interaction network and functional enrichment analysis. Genomics Inform 18, e39 (2020). 10.5808/GI.2020.18.4.e39

40 Puar, N., Chovatiya, R. & Paller, A. S. New treatments in atopic dermatitis. Ann Allergy Asthma Immunol 126, 21–31 (2021). 10.1016/j.anai.2020.08.016

41 Frew, J. W. et al. A Systematic Review of Promising Therapeutic Targets in Hidradenitis Suppurativa: A Critical Evaluation of Mechanistic and Clinical Relevance. J Invest Dermatol 141, 316–324 e312 (2021). 10.1016/j.jid.2020.06.019

42 Kim, J. et al. Single-cell transcriptomics suggest distinct upstream drivers of IL-17A/F in hidradenitis versus psoriasis. J Allergy Clin Immunol 152, 656–666 (2023). 10.1016/j.jaci.2023.05.012

43 Mariottoni, P. et al. Single-Cell RNA Sequencing Reveals Cellular and Transcriptional Changes Associated With M1 Macrophage Polarization in Hidradenitis Suppurativa. Front Med (Lausanne) 8, 665873 (2021). 10.3389/fmed.2021.665873

44 Gudjonsson, J. E. et al. Contribution of plasma cells and B cells to hidradenitis suppurativa pathogenesis. JCI Insight 5 (2020). 10.1172/jci.insight.139930

45 Biajoux, V. et al. Efficient Plasma Cell Differentiation and Trafficking Require Cxcr4 Desensitization. Cell Rep 17, 193–205 (2016). 10.1016/j.celrep.2016.08.068

46 Miller, R. J., Banisadr, G. & Bhattacharyya, B. J. CXCR4 signaling in the regulation of stem cell migration and development. J Neuroimmunol 198, 31–38 (2008). 10.1016/j.jneuroim.2008.04.008

