## Supplementary figures and images for "Integrative systems biology framework discovers common gene regulatory signatures in multiple mechanistically distinct inflammatory skin diseases"

### Supplemental Figure 1

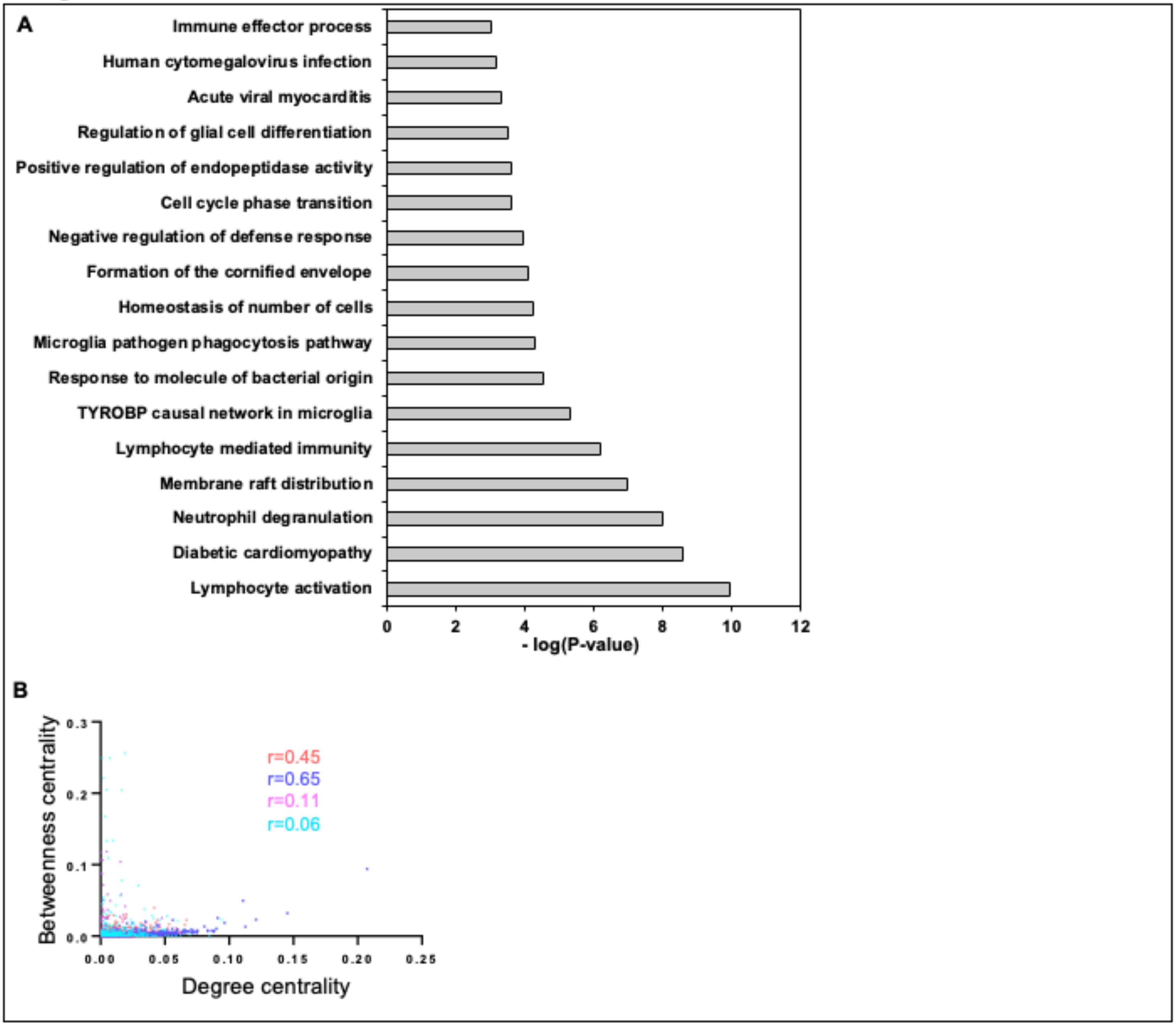
